# High-plex Digital Spatial Profiling Identifies Subregion-Dependent Directed Proteome Changes Across Multiple Variants of Dementia

**DOI:** 10.1101/2024.04.29.591721

**Authors:** MacKenzie L. Bolen, Kelly B. Menees, Andrea R. Merchak, Marla Gearing, Jingjing Gong, Yuqi Ren, Melissa Murray, Zachary T. McEachin, Malú Gámez Tansey

**Author notes:** **Correspondence:** Dr. Malú Gámez Tansey, Department of Neuroscience, University of Florida, College of Medicine, Gainesville, Fl, USA. **Contributor information:** MacKenzie Bolen, Kelly B. Menees, Andrea R. Merchak, Marla Gearing, Jingjing Gong, Yuqi Ren, Melissa Murray Zachary T McEachin.

## Abstract

Frontotemporal lobar degeneration (FTLD) is the leading cause of dementia in patients under the age of 65. Even in a single anatomical region, there is variance within pathological protein deposition as a result of FTLD. This spectrum of pathology leads to difficulty in identification of the disease during its progression and consequential varied post mortem clinicopathological diagnoses. NanoString GeoMx™ Digital Spatial Profiling (DSP) is a novel method that leverages the ability to spatially multiplex protein biomarkers of interest. We utilized NanoString DSP to investigate the proteome geography at two levels of the cortex and the subcortical white matter in patients with various types of dementia (Alzheimer’s disease, FTLD-c9ALS, FTLD-17, FTLD-TDP, FTLD-GRN; n=6 per syndrome) and neurologically healthy controls (NHC). Analysis of 75 different protein biomarkers of interest revealed both a disease and cortical subregion specific biomarker profile. Layers II-V of the cortex from diseased individuals displayed the greatest protein dysregulation as compared to NHC. Additionally, out of all disease groups within cortical layer 1, II-V and white matter, the FTLD-17 group had the most significant protein dysregulation as compared to NHC--specifically associated with immune cell pathways. Traditional biomarkers of dementia, such as various phosphorylated tau proteins and amyloid-β 42 displayed dysregulation, however our data suggest spatial enrichment in distinct to cortical sublayers. In conclusion, we observed that depending on the variant of disease, specific protein deposits either span multiple levels of cortical geography or only a single layer. Thus, the specific localization of these deposits could potentially be used to elucidate region-specific pathologic biomarkers unique to individual variants of dementia.

## INTRODUCTION

FTLD encapsulates a group of degenerative proteinopathies associated with the early breakdown of the frontal and temporal lobes of the brain and is the neuropathological diagnosis corresponding to clinical frontotemporal dementia (FTD) [1, 2]. Early stages of disease are commonly characterized by language difficulties and behavioral abnormalities, while later stages of FTLD constitute a rapid cognitive decline paired with dementia [1, 3].There are three primary clinical groupings of FTD: the behavioral variant (bvFTD), progressive non-fluent aphasia, and movement semantic dementia [1, 4]. However, even within these groupings there is vast non-linear heterogeneity across disease pathology, etiology and symptomology [5], which has presented significant challenges to therapeutic research and pathological diagnosis within the FTLD spectrum.

Up to 40% of all FTLD cases are associated with various genetic variants [4,5]. The most common genetic variants associated with FTLD are linked to changes in the microtubule associated protein tau *(MAPT*), progranulin (*PGRN*), chromosome 9 open reading frame 72 (*C9ORF72*) and transactive DNA-binding protein gene (*TARDBP*) that expresses TDP-43 [6].

Interestingly, FTD presentation and prognosis can overlap to some extent with Alzheimer’s disease (AD), Parkinson’s disease (PD), and amyotrophic lateral sclerosis (ALS) – making it difficult to distinguish FTD from AD, PD, or ALS until the pathological diagnosis can be determined [7,8,9]. Making the diagnosis even more complicated is comorbid disease onset; meaning a patient can clinically present with FTD but the pathological diagnosis at autopsy may include secondary pathology such as AD neuropathologic changes, Lewy bodies, or TDP-43-positive inclusions.

A newly emerging thread of investigation to better diagnose FTLD and related dementias postmortem has been region specific analysis of protein deposition. FTLD appears to induce syndrome specific protein deposition [10,11]; however, the distinct protein community(s) unique to each cortical subregion has yet to be characterized. The cortical layers each have unique cellular compositions, resulting in a diverse array of complex functions. With the understanding that the catalyst of FTLD can be patient specific and extremely variable, the current frontier of FTLD research has moved towards a more multiplexed approach to investigate the role laminar cortical pathology plays in the clinical syndrome and proteinopathies associated with FTLD and related dementias.

In this study, we aimed to identify differentially abundant proteins in a spatially resolved manner across several neurodegenerative diseases including AD and several FTLD subtypes. Here, we utilize NanoString GeoMx DSP, a novel method that leverages the ability to spatially multiplex a community of protein biomarkers of interest. This technology provides a single system approach to investigating the region specific pathologic proteomic contribution of specific disease states, which will lay the groundwork for future biomarker investigation and clarification of the FTLD disease spectrum.

## MATERIALS AND METHODS

### Cases

Cortical tissue was obtained from Emory University Goizueta Alzheimer’s Disease Research Center (ADRC) brain bank on 30 patients diagnosed with neurological disease and 6 neurologically healthy controls (Table 1: Additional File 1). The five neurological diseases assessed include Alzheimer’s disease (AD), amyotrophic lateral sclerosis associated with a C9 expansion mutation (C9ALS), frontotemporal lobar degeneration with parkinsonism-17 (FTDP-17), frontotemporal lobar degeneration with TPD-43-immunoreactive pathology (FTLD-TDP), and progranulin mutation related FTLD-TDP (GRN). Disease and neurologically healthy control groups were matched as closely as possible for age and sex. Neuropathologic diagnosis and characterization of disease associated pathology was conducted at Emory University (MG) following standardized criteria [12,13]. Individual case Braak staging and secondary neuropathology are detailed in Supplementary Table 5.

**Table 1.**
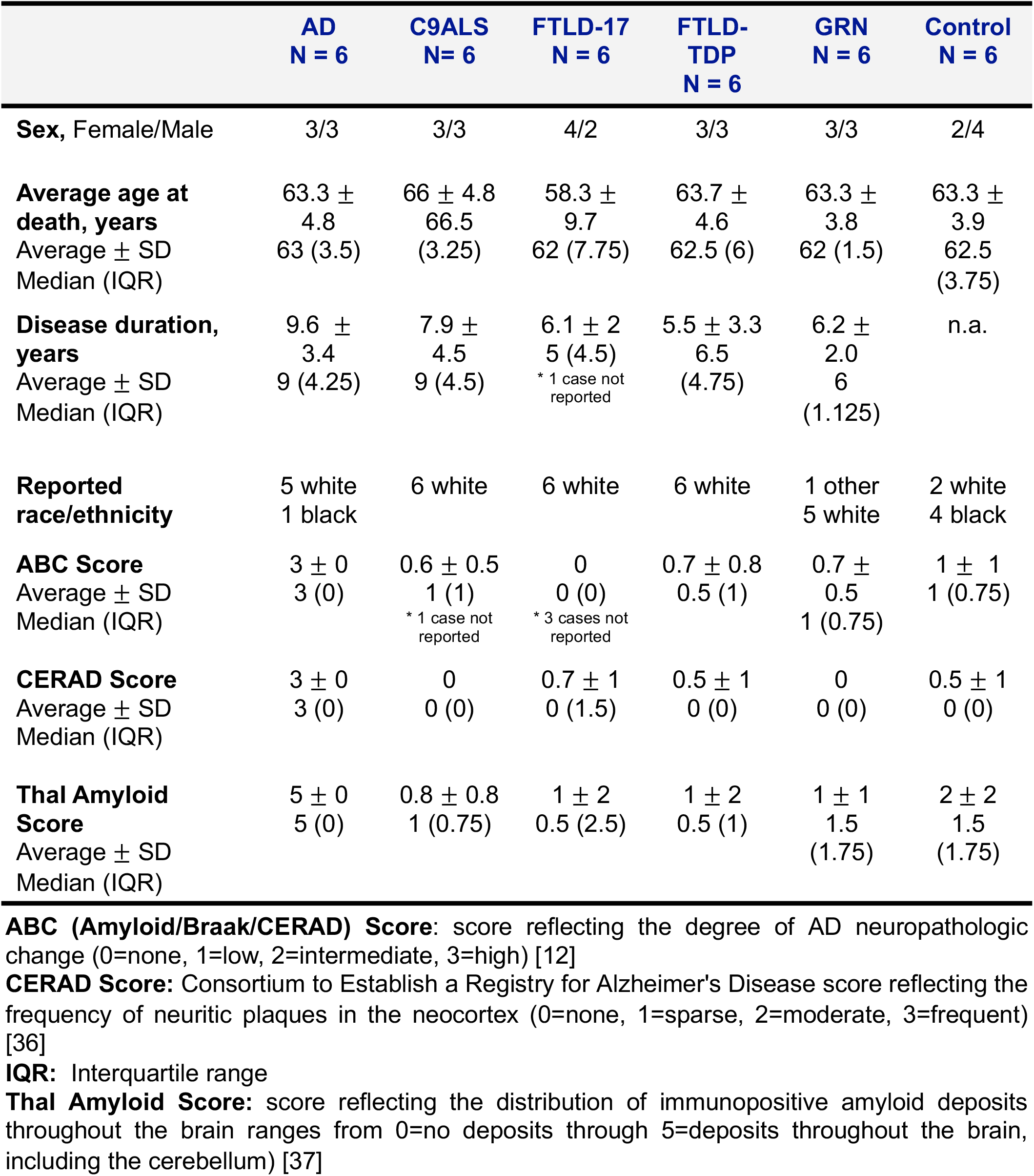
Case demographics, cognitive status and neuropathologic outcomes (1.75) (1.75)

### NanoString GeoMx™ Digital Spatial Profiling (DSP)

Paraformaldehyde-fixed prefrontal cortex samples were paraffin embedded and sectioned at 5μm. Sections were then deparaffinized and incubated with the NanoString GeoMx Human Neuroscience panel (Fig. 1C); which includes 75 UV-cleavable protein probes and 3 antibody visualization markers against nuclear DNA (Syto13), microglia (IBA1, Cat#MABN92-AF647) and astrocytes (GFAP, Cat#8152S) (Fig. 1A and visualization in Fig. 1B). Tissue rehydration and antibody incubation was completed according to the manufacturer’s instructions. Visualization markers were used to select regions of interest (ROI). ROIs were defined by cortical location, being molecular (layer I), internal layers (layers II-V) and subcortical white matter. Cortical layers were differentiated by density of nuclear staining and geographic location relative to tissue edge. Twelve ROIs per case were chosen, where four ROIs were drawn per cortical layer (Fig. 1B). Once ROIs were selected on the GeoMx™ digital spatial profiler, UV light was used to illuminate the ROI – cleaving the protein specific UV-cleavable oligonucleotide tags. ROI specific cleaved tags then hybridized overnight to unique to six-color optical barcodes. Optical barcodes were then quantified on the NanoString nCounter MAX/FLEX by counting the number of barcodes specific to each protein present within each ROI. All digital counts were normalized to the geometric mean of the ROI area.

**Fig. 1.**
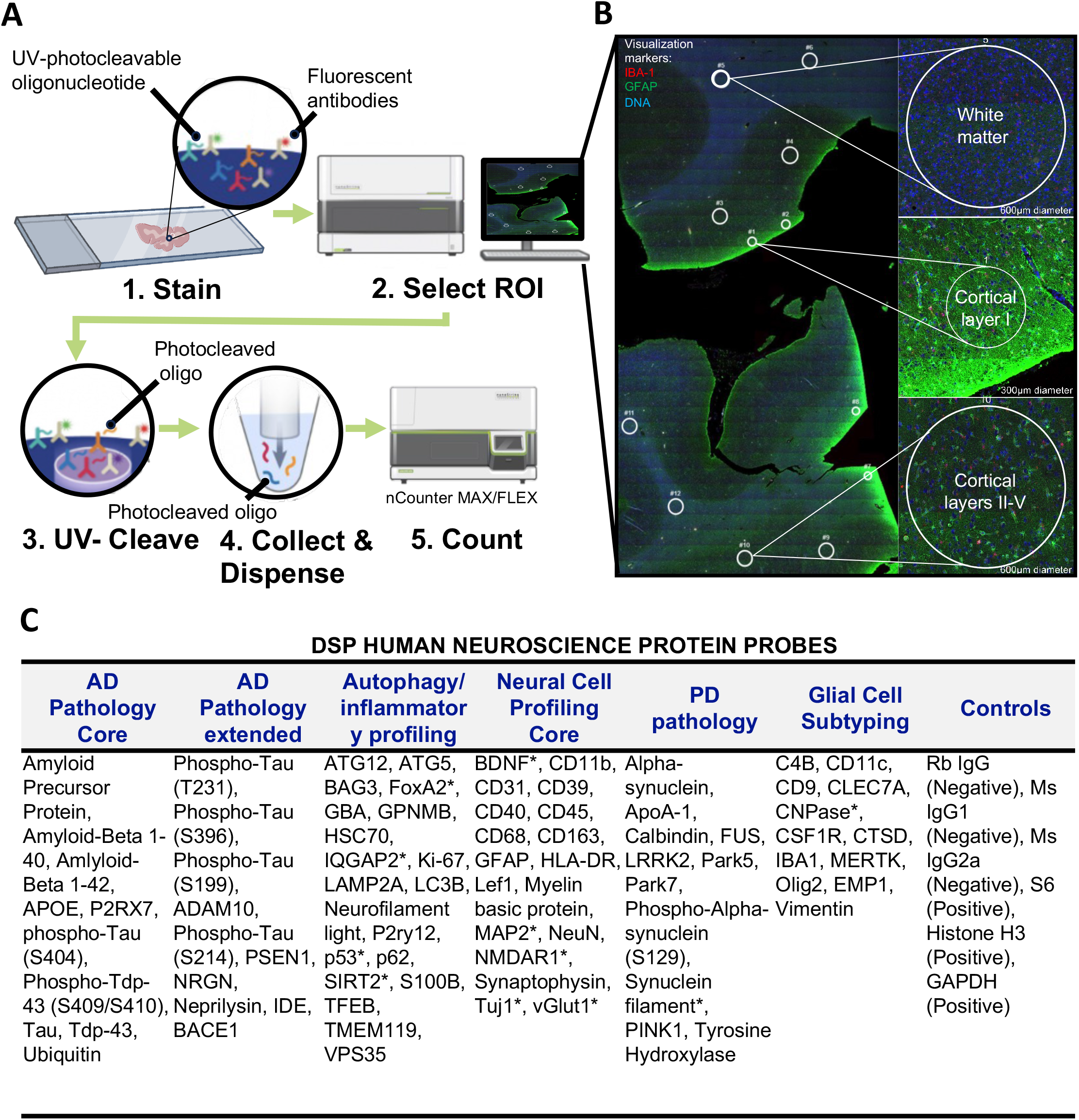
Digital spatial proteomic profiling (DSP) using immunofluorescence as an indicator of multiple proteomic targets on a single slide. **A**. Working procedure of NanoString GeoMx Digital Spatial Profiling using nCounter MAX/FLEX. Formalin-fixed paraffin-embedded cortical human tissue is incubated with 3 antibody visualization markers, followed by incubation with 75 DSP probes. After tissue is imaged, regions of interest (ROI) are digitally selected, ROIs are then subjected to ultraviolet (UV) light which results in release of UV-cleavable protein specific oligonucleotide barcodes from DSP probes. Cleaved protein unique barcodes are then collected and quantified. **B**. Microphotograph of cortical tissue labeled with Iba1 (red), GFAP (green) and DNA (blue) visualization markers. Regions of interest (ROI) were manually drawn in approximately the same location within white matter, molecular layer (cortical layer I) and internal/granular layer (cortical layer IV) for every case. Four ROIs per subregion per patient (12 ROIs per tissue section) were collected and assessed as individual data points in all downstream analyses. **C**. DSP human Neuroscience protein probes, separated by disease or functional genre, a (*) indicates a protein that was not part of the NanoString Human Neuroscience panel but rather was chosen by our lab and spiked in.

### GeoMx™ DSP analysis

Raw protein expression data was input into NanoString DSPAPP V6.0 Analysis Software. Expression of 75 different proteins was analyzed using the Human Neuroscience panel specific endogenous controls, which act as an internal quality check for technical variation. Then counts were normalized to surface area (geometric mean) of each individual ROI and the expression of negative/positive and housekeeping controls (GAPDH, Histone H3, S6). ROIs within each cortical layer of each case were then compared to other cortical layers across disease type. Differential expression analysis was completed via a linear mixed model, adjusted for age and sex. P-values were adjusted for multiple analyses via Benjamini-Hochberg procedure. Heatmaps displaying p-values were created by exporting the normalized .csv File from GeoMx ™ analysis suite and importing into GraphPad Prism (Version 10.1.1).

## RESULTS

### Cortical sublayers reveal disease-relevant shared and distinct protein expression

By employing the NanoString GeoMx DSP platform we assessed the regulation of 75 different proteins across 3 separate layers (molecular layer I, internal layers II-V and white matter) of cortical tissue, in 5 different types of dementia (Fig. 1A,B,C). Individual cortical subregion ROI analysis of the cortex revealed differential disease associated protein regulation (Fig. 2). Differential expression was fit on a per protein basis using a linear mixed model. The model used the protein expression as the dependent variable and fit the cortical subregions and diseases as a fixed effect respectively to test for differences between groups. Patient was fit as a random effect. We grouped these protein communities by NanoString recommended panel association or by autophagy and/or inflammation associated proteins (Fig. 1C), the composition of proteins drastically changes within each community based off cortical subregion. We were able to parse apart known disease relevant biomarkers and communities of implicated biomarkers that are subregion dependent. For example, amyloid beta 42 (Aβ-42) and phosphorylated tau S214, S396 and T231 deposition only significantly increased in the molecular and internal sublayers of the cortical tissue from AD patients assessed (Fig. 2B,C) but not the subcortical white matter (Fig. 2A). Interestingly, there was a significant decrease in all phosphorylated tau in subcortical white matter tissue in the AD, c9ALS, FTLD, and GRN pathology groups (Fig. 2A) but a significant increase of phosphorylated tau in the internal and molecular layers of the cortex in AD and FTLD-17 tissue, as compared to neurologically healthy control (Fig. 2B,C). When comparing significantly different protein expression in disease groups vs NHC, no disease group showed significant overlap in protein expression across cortical subregion (Fig. 2).

**Fig. 2.**
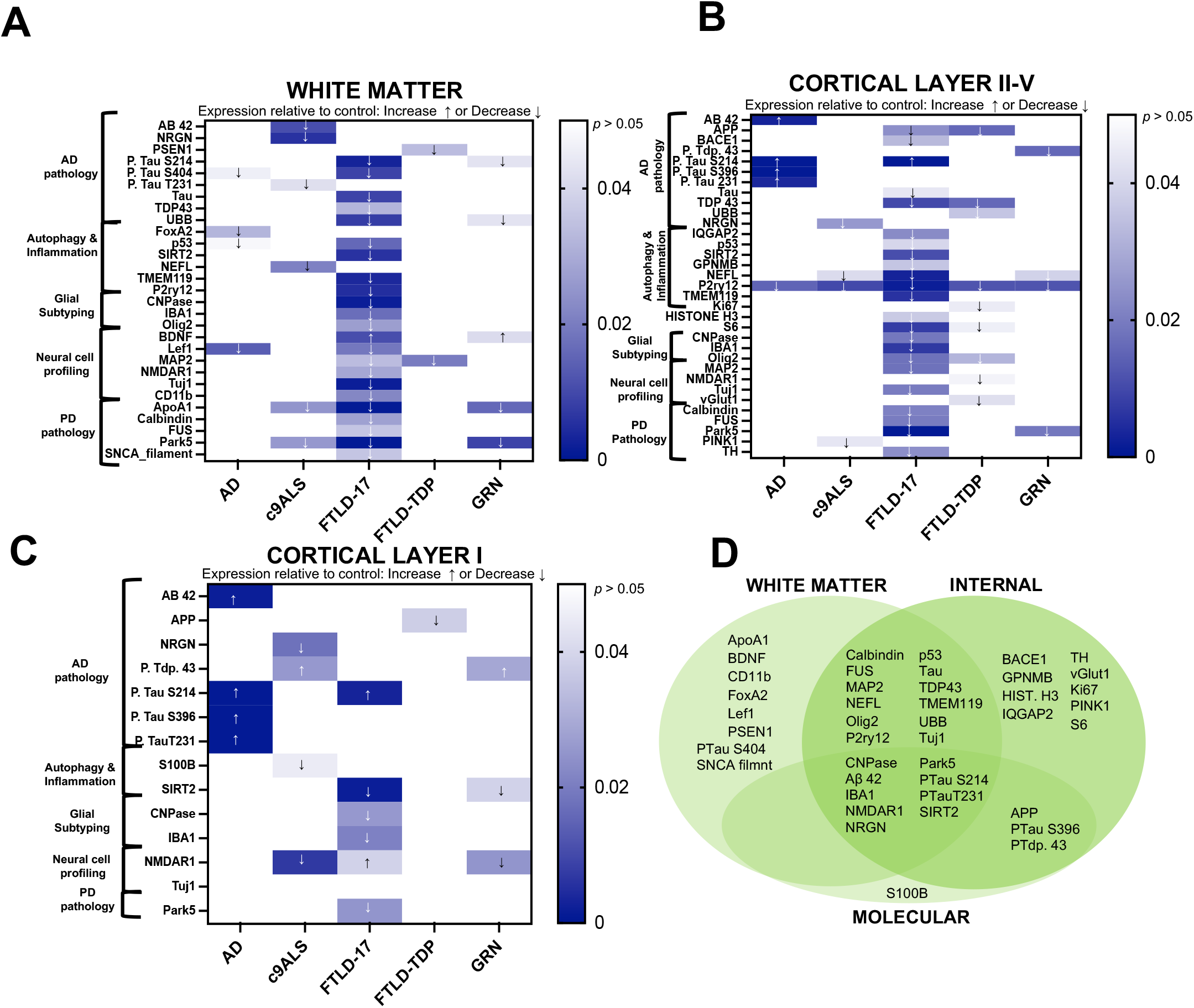
Cortical and subcortical laminar layers reveal significantly different disease specific proteomic expression relative to control. **A-C** Heatmaps of differentially expressed proteins when comparing all cortical subregions of interest across disease diagnosis. Linear mixed model regression analysis was used to identify significantly upregulated proteins in disease groups are indicated as any value above zero but less than *p* < 0.05. Significantly (*p* < 0.05) downregulated proteins are indicated by any value below zero. Individual significance and fold change values by disease within each subregion can be found in Supplemental Figures 2-5. **D**. Venn diagram depicting proteins distinct to cortical subregions within all disease groups that are significantly different than control. All counts were normalized to ROI area and internal controls.

### Spatial profiling of cortical white matter reveals significant protein downregulation in 5 variants of dementia across multiple cell types, compared to neurologically healthy control

Proteins downregulated across multiple diseases in subcortical white matter include Park-5 (c9ALS, FTLD-17, GRN), ApoA1 (c9ALS, FTLD-17, GRN), Map2 (FTLD-17 and FTLD-TDP) phosphorylated tau S214 (FTLD-17 and GRN), UBB (FTLD-17 and GRN), p53 (AD and FTLD-17), and Lef1 (AD and FTLD-17) (Fig. 2A: Additional File 1). The only upregulated protein across multiple diseases in the white matter was BDNF (FTLD-17 and GRN) (Fig. 2A). The FLTD-17 positive cohort had the lowest expression of all significantly different proteins in the subcortical white matter as compared to AD, c9ALS, FTLD-TDP and GRN (Fig. 2A: Additional File 1).

Collectively these data suggest that independent of disease, some protein deposition indicative of pathologic progression may be unique to specific cortical layers.

### Glial activation marker, P2ry12, is downregulated across all five variants of dementia in the internal cortical laminar layer

P2ry12 was significantly downregulated across all five disease groups, as compared to NHC in the internal subregion of the cortex (Fig. 2B: Additional File 2), there was no other protein within our chosen panel that impacted all 5 disease groups within the same region. Proteins downregulated across multiple groups within the internal subregion of the cortex include APP (significant change identified in FTLD-17 and FTLD-TDP), TDP-43 (significant change identified in FTLD-17 and FTLD-TDP), NEFL (significant change identified in c9ALS, FTLD-17 and GRN), S6 (significant change identified in FTLD-17 and FTLD-TDP), Olig2 (significant change identified in FTLD-17 and FTLD-TDP) and Park-5 (significant change identified in FTLD-17 and GRN) (Fig. 2B: Additional File 2). The only protein upregulated across multiple diseases within the internal sublayer of the cortical tissue assessed was phosphorylated tau S214 (significant change identified in FTLD-17 and GRN) (Fig. 2B: Additional File 2). These data suggest that multiplexing P2ry12 with pathological indicators of disease, like TDP-43, NEF-L and phosphorylated tau S214, could provide additional insight into parsing apart the discrepancies between dementia variant associated protein dysregulation and deposition.

### In the molecular laminar layer of the cortex, phosphorylated tau S214 and/or TDP-43 are upregulated in various forms of dementia but not all

Within the molecular layer of the cortex, NMDAR1 was downregulated in both C9ALS and GRN cortical tissue, the protein was upregulated in FTLD-17 (Fig. 2C: Additional File 3). The proteins upregulated across multiple neurodegenerative diseases within the molecular layer include phosphorylated TDP-43 (significant change identified in c9ALS and GRN) and phosphorylated Tau S214 (significant change identified in AD and FTLD-17) (Fig. 2C: Additional File 3). Phosphorylated TDP-43 and phosphorylated tau S214 are two of many pathologic lesions associated with FTLD [15], of interest is the region-specific upregulation of these markers increase in deposition that appears to be disease dependent within the FTLD spectrum. For example, phosphorylated tau S214 is upregulated in both the internal and molecular cortical layers but down regulated in the white matter of cortical tissue positive for FTLD-17. These data reveal distinct spatial distribution of known pathology associated proteins in a disease dependent fashion.

### Pathologic protein deposits develop within specific brain regions independent of disease variant

Using BioVenn, we created a Venn diagram depicting the grouped relationships of protein expression by region, independent of disease group (Fig. 2D). Interestingly, the molecular subregion of the cortical tissue had the least amount of significant protein differences when comparing disease groups to NHC, where only 4% of all significant proteins were exclusively expressed within the molecular sublayer (Fig. 2D). The white matter and internal sublayers of the cortex shared 26% of the total significant protein change. Additionally, 21% of all proteins were significantly different when comparing any disease group vs NHC in all three cortical subregions assessed (Fig. 2D). Collectively these data suggest that individual cortical and subcortical layers may be more likely to reveal patterns of protein deposition than others.

### Amyloid and dopaminergic protein biomarkers transcend cortical subregion specificity across multiple variants of dementia

The final analysis compared all ROIs (n = 4 per layer) collected across all cortical sublayers from every patient, where all disease ROIs were compared against each other and NHC (Fig. 3). As anticipated a handful of canonical biomarkers of neurodegenerative pathology, such as Aβ-42, NMDAR1, and phosphorylated tau T231, revealed differential regulation as compared to NHC independent of ROI subregion (Fig. 3 A-D: Additional File 4). On the contrary, other popular canonical markers of neurodegeneration associated with dementia appear to be subregion specific. For example, amyloid precursor protein (APP), a protein commonly associated with dementia, only indicated significant downregulation relative to NHC in the molecular and internal subregions of the cortex but not the subcortical white matter (Fig. 2: Additional Files 1-3). Of additional interest was tyrosine hydroxylase, an indicator of dopamine synthesis that is commonly decreased in parkinsonism associated dementia, was only found to significantly decrease from NHC in the internal sublayer of the cortex with no significant change in expression in any other investigated ROI (Fig. 3: Additional File 4). These data suggest that not all protein regulation (observed from the chosen panel used within this study), is localized to a single layer. In the context of the 5 dementia variants investigated, some protein regulation may be domestic to each cortical layer, while other disease relevant proteins reveal a global response across the cortex.

**Fig. 3.**
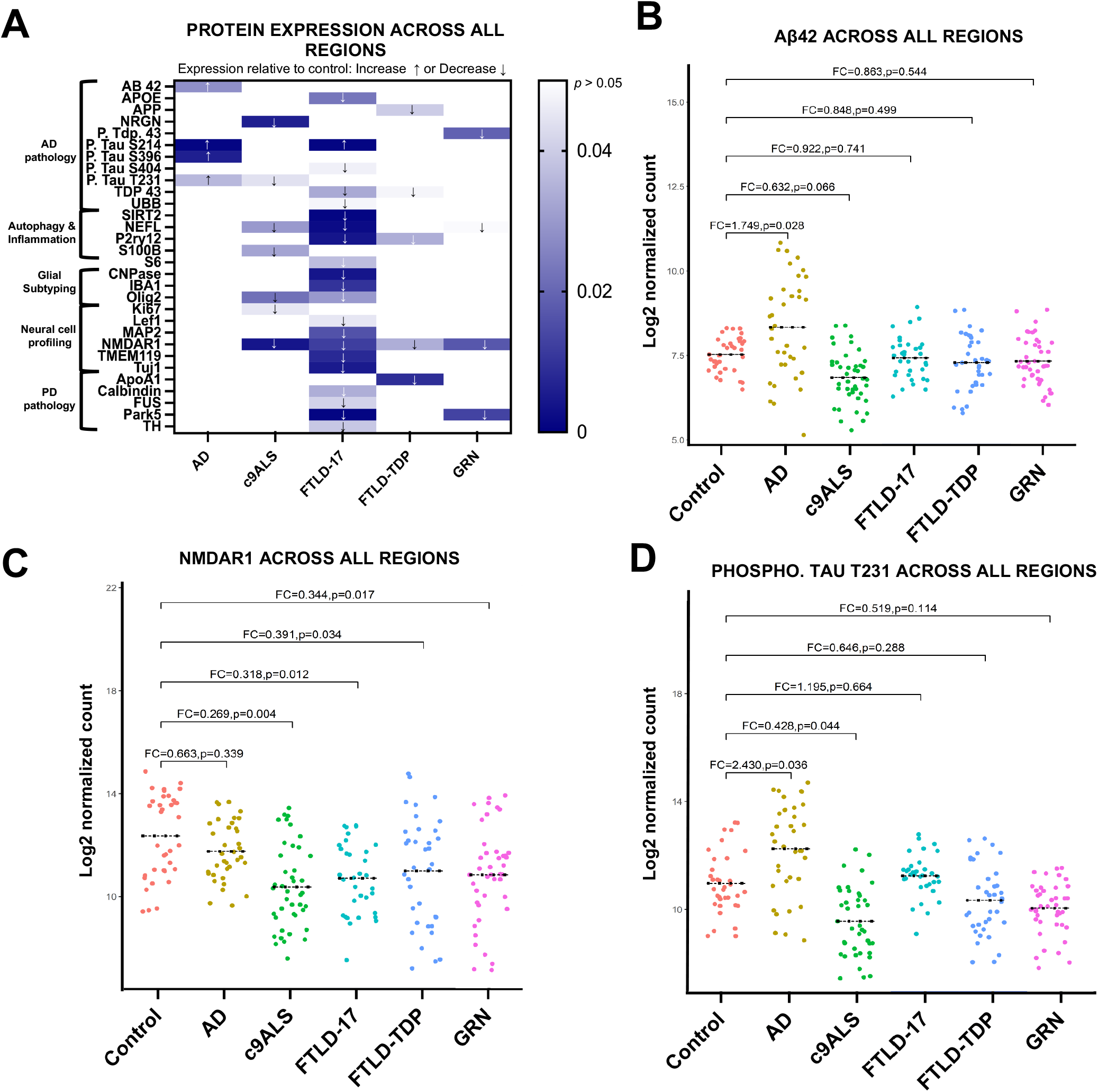
DSP identifies significant disease dependent protein expression changes irrespective of laminar cortical and subcortical layers of the brain. **A**. Heatmap of differentially regulated proteins when comparing all regions of interest. Linear mixed model regression analysis was used to identify significantly upregulated proteins disease groups are indicated as any value above zero but less than *p* < 0.05. Significantly (*p* < 0.05) downregulated proteins are indicated by any value below zero. **B-D** Individual protein markers of interest independent of subregion specificity (all ROIs merged). Generalized linear regression comparing Log2 normalized gene expression via quantification of oligonucleotide barcode reads (counts) between disease groups was considered significant when *p* > 0.05. All counts were normalized to ROI area and internal controls.

## DISCUSSION

FTLD is characterized by the progressive breakdown of both the anterior frontal and temporal lobes of the brain [1,2] and is one of the most common causes of early onset dementia. Within the web of FTLD, there is vast heterogeneity in pathologic protein deposition in both the frontal and temporal cortices, resulting in a spectrum of clinical symptomology and pathology [16,17,18,19,20,21]. Therefore, the clinicopathologic diagnosis of FTLD and associated comorbidities can be extremely challenging [22], and the pathogenic mechanisms underlying disease onset and progression continue to be elusive.

The cortex is highly specialized, with each layer serving a distinct role. It remains unclear as to what drives unique subregion dependent protein deposition due to disease. By leveraging the NanoString GeoMx DSP technology we were able to demonstrate the extensive proteomic differences induced by disease and guided by cortical and subcortical laminar region. Not only did we further validate Aβ-42 and phosphorylated tau regulation modulation in the cortex and subcortical white matter layer as a result of disease, but we more distinctly parsed which communities of protein biomarkers also changed in conjunction with these canonical pathologic indicators. We identified distinct communities of proteins that were unique to each cortical subregion, elucidating a critical role spatial proteomics can play in further characterizing the complex mechanisms of neurodegenerative disease.

In addition to further validation of canonical biomarkers of FTLD, we were able to distinguish phosphorylation patterns of pathological proteins (like tau) that were unique to each cortical subregion. These data reveal that phosphorylated tau protein deposits are likely unique to cortical and subcortical subregion. Although there is substantial evidence of various phosphorylated tau protein deposition in multiple regions of the brains of FTLD patients [23,24,25], our data further solidify that phosphorylated tau may be subregion dependent within the human cortical and subcortical brain layers.

Of interest are the roles that inflammation and the immune system may play in subsequent brain region-specific protein deposition. P2ry12, a marker associated with broad microglia activation, significantly decreased in the subcortical white matter of only FTLD-17 patients and also decreased within layers II-V of the cortex in AD, c9ALS, FTLD-17, FTLD-TDP and GRN patients (Fig. 2). Iba1, another general marker for activated microglia, is significantly downregulated in FTLD-17 patients within the white matter, layer I and layers II-V, as compared to NHC (Fig. 2). Depletion in Iba1 positive microglia in the human brain has been reported previously in association with both early- and late-onset AD [26,27,28] however within our data set we only observe this phenotype in FTLD-17. Additionally, we identified a significant depletion in Sirt2 within the white matter, layer I and layers II-V, as compared to NHC (Fig. 2). Depletion of Sirt2 may be an indicator of mitochondrial stress and resulting apoptosis [29]. CNPase, a marker for myelin integrity and maintenance [30], was significantly downregulated in FTLD-17 patients as compared to NHC within the white matter, layer I and layers II-V, (Fig. 2). A decrease in p2ry12 and Iba1-positive microglia paired with dysfunctional myelin machinery and an increase in apoptotic features in FTLD-17 patients suggests multi-level cortical dysregulation of immune activity and resilience. These data suggest that FTLD-17 induces the largest proteomic dysregulation associated with inflammatory distress in white matter and cortical layers I, II-V, as compared to what we detected in FTLD, AD and NHC samples.

Although our data support distinct cortical subregion proteomic profiles, our analysis also indicates that a handful of mislocalized proteins transcend cortical layer specificity (Fig. 3). As expected, increased deposition of Aβ42 was evident in AD tissue when all regional ROIs were compared against NHC (Fig. 3). Interestingly NMDAR1 significantly decreased in c9ALS, FTLD-17, FTLD-TDP and GRN tissue relative to NHC when all ROIs were combined (Fig. 3). NMDAR1 protein expression has been reported to decrease due to glutamatergic synaptic depression in tissue burdened with either tau [31] or Aβ pathology [32]. Our data reveal that NMDAR1 may represent a viable biomarker for multiple variants of FTLD but not AD (Fig. 3).

Previous proteomic studies investigating FTLD or related dementia in human brains have assessed regional and/or cell-specific protein content [22,33,34]. NanoString DSP has been successfully used to delineate region-specific immune microenvironments associated with brain samples from individuals pathologically diagnosed with the TREM2 risk variant for AD [38] and in separate study, parse apart neuroprotective protein regulation in brain samples from individuals resilient to AD cognitive consequence [43]. However, no study has reported multiplexed detection of different pathologic proteins of interest within each subregion of the cortex across multiple syndromes of FTLD. Common techniques studying protein-associated pathology such as mass spectrometry and western blotting are still highly relevant and used widely in the field. NanoString DSP does not replace these techniques but supplements these well-established tools by enabling preservation of the spatio-anatomical localization of proteins of interest. For example, in induced pluripotent stem cell (iPSC)-derived neurons from patients with various FTLD mutations, a dysregulated relationship between tau and mitochondrial proteins was identified using ascorbic acid peroxidase (APEX) combined with quantitative affinity purification mass spectrometry (AP-MS) [35]. Although the authors were able to provide insight on structural morphology by imaging the neurons, these data do not provide insight into the relevance of anatomical location within the complex and heterogenous microenvironment of the brain. Additional recent reports have provided region-specific insight into the early PD proteome, where 8 brain regions were spatially investigated using MS^e^ label free proteomics [34]. Although this method provides excellent region-specific protein interactions, the data set is specific to early PD and does not provide insight into subregion specific protein regulation.

Although our findings are novel and significant, the studies herein suffer from several limitations that should be noted. GAPDH, Histone H3, and S6 were used as internal controls to normalize our data set, although there was a significant decrease in Histone H3 and S6 in the internal layer of FTLD-TDP patients, as compared to NHC. We acknowledge that this outcome indicates that Histone H3 and S6 are not ideal quality controls for the internal cortical layer of FTLD-TDP patients, which further indicates the need for not only cortical subregion-dependent investigation but also subregion-specific downstream normalization. Additionally, the number of cases per disease group was relatively small (n=6), with such high variation within said small groups we chose to treat each ROI within each subregion as its own representative data point. Future studies should consider the variation within each patient and use these data for determination of ROI-independent statistical assessment. These data help build a basis for region-specific proteomic assessment of FTLD and provide diagnostic insight into the region-specific discrepancies between each syndrome. Transcriptome information and single-cell resolution could further parse mechanistic differences (such as providing insight into specific cell-type ligand-receptor interactions and cell-type neighborhood interactomes) between various FTLD syndromes. Future work using single cell spatial omics technology, like 10X Genomics or NanoString CosMx SMI, would provide additional insight into this critical line of research.

## CONCLUSIONS

NanoString GeoMx™ is a novel method that leverages the ability to spatially multiplex protein biomarkers of interest. With the understanding that the catalyst of FTLD can be patient specific and extremely variable [39,40,41,42], the current frontier of FTLD and neurodegeneration research has moved towards spatial proteomics approaches, where focused brain region-specific assessments now enable investigators to gain additional insight into the pathologic mechanisms underlying this spectrum of neurodegenerative syndromes. Our data provide a unique perspective on subregion-specific protein expression and/or depositions in various syndromes of FTLD that traditional proteomic approaches are unable to capture with precision. Specifically, the FTLD-17 spatial proteome appears to be enriched with the largest immune- and inflammation-related dysregulation across both gray and white matter in the cortex. By spatially mapping protein expression within the cortex of five different variants of dementia we were able to catalogue and continue to parse apart the brain region- and disease-specific proteome of various dementia syndromes. While a bit surprising to us, we find that although some traditional histopathologic markers of interest to dementia researchers selectively localize to specific brain regions of the cortex, other potentially pathologic protein deposits span wider anatomical areas of the brain. These data lay a foundation for additional work aimed at parsing out common and shared mechanisms underlying various forms of FTLD disease in variant-specific future biomarker discovery efforts.

## Acknowledgements

We would like to thank the Tansey lab for critical feedback on these data. We would also like to thank all of the participants who donated tissue to the Goizueta Alzheimer’s Disease Research Center at Emory University and its Director Dr. Allan I. Levey, without them this work would not have been possible. Figure 1A was created partially by BioRender.com.

## ABREVIATIONS

AD: Alzheimer’s disease
Aβ: Amyloid-β
ABC score: Activities of daily Living, Behavioral and psychological Symptoms of Dementia, and Cognitive Function
AP-MS: quantitative affinity purification mass spectrometry
APP: Amyloid precursor protein
APEX: Ascorbic acid peroxidase
ApoA1: Apolipoprotein A1
ALS: Amyotrophic lateral sclerosis
bvFTD: Behavioral variant frontotemporal dementia
CERAD score: Consortium to Establish a Registry for Alzheimer’s Disease
C9ORF72: chromosome 9 open reading frame 72
DSP: Digital spatial profiling
FDR: False discovery rate
FTD: Frontotemporal dementia
FTLD: Frontotemporal lobar degeneration
GAPDH: glyceraldehyde-3-phosphate dehydrogenase
GFAP: Glial fibrillary acidic protein
GRN: progranulin mutation related frontotemporal lobar degeneration
IBA1: Ionized calcium binding adaptor molecule 1
IPSC: Induced pluripotent stem cell
IQR: Interquartile range
Lef1: Lymphoid Enhancer Binding Factor 1
MAPT: microtubule associated protein tau
Map2: Microtubule associated protein 2
NEFL: Neurofilament light
NHC: Neurologically healthy control
NMDAR1: Glutamate [NMDA] receptor subunit 1
Park5 (UCH-L1): Ubiquitin carboxyl-terminal hydrolase isozyme L1
PD: Parkinson’s disease
PGRN: Progranulin
P2ry12: Purinergic receptor P2Y12
P53: Tumor protein 53
ROI: Region of interest
S6: Ribosomal protein S6
TARDBP: Transactive DNA-binding protein gene
TDP-43: Transactive response DNA binding protein of 43 kDa
UBB: Ubiquitin

## REFERENCES

1. Rabinovici GD, Miller BL (2010) Frontotemporal lobar degeneration: epidemiology, pathophysiology, diagnosis and management. CNS Drugs 24:375–398. 10.2165/11533100-000000000-00000

2. Murray ME, DeTure M (2018) The neuropathology of dementia. In: Smith GE, Farias ST (eds) APA handbook of dementia. American Psychological Association, Washington, pp 41–66

3. Snowden J, Neary D, Mann D (2007) Frontotemporal lobar degeneration: clinical and pathological relationships. Acta Neuropathol 114:31–38. 10.1007/s00401-007-0236-3

4. Hodges JR, Miller B (2001) The classification, genetics and neuropathology of frontotemporal dementia. Introduction to the special topic papers: Part I. Neurocase 7:31–35. 10.1093/neucas/7.1.31

5. Bocchetta M, Malpetti M, Todd EG, et al (2021) Looking beneath the surface: the importance of subcortical structures in frontotemporal dementia. Brain Commun 3:fcab158. 10.1093/braincomms/fcab158

6. Goldman JS, Farmer JM, Wood EM, et al (2005) Comparison of family histories in FTLD subtypes and related tauopathies. Neurology 65:1817–1819. 10.1212/01.wnl.0000187068.92184.63

7. Alladi S, Xuereb J, Bak T, et al (2007) Focal cortical presentations of Alzheimer’s disease. Brain 130:2636–2645. 10.1093/brain/awm213

8. Womack KB, Diaz-Arrastia R, Aizenstein HJ, et al (2011) Temporoparietal hypometabolism in frontotemporal lobar degeneration and associated imaging diagnostic errors. Arch Neurol 68:329–337. 10.1001/archneurol.2010.295

9. Pasquier F (2005) Telling the difference between frontotemporal dementia and Alzheimer’s disease. Curr Opin Psychiatry 18:628–632. 10.1097/01.yco.0000185988.05741.2a

10. Ohm DT, Cousins KAQ, Xie SX, et al (2022) Signature laminar distributions of pathology in frontotemporal lobar degeneration. Acta Neuropathol 143:363–382. 10.1007/s00401-021-02402-3

11. Vatsavayai SC, Nana AL, Yokoyama JS, Seeley WW (2019) C9orf72-FTD/ALS pathogenesis: evidence from human neuropathological studies. Acta Neuropathol 137:1–26. 10.1007/s00401-018-1921-0

12. Montine TJ, Phelps CH, Beach TG, et al (2012) National Institute on Aging-Alzheimer’s Association guidelines for the neuropathologic assessment of Alzheimer’s disease: a practical approach. Acta Neuropathol 123:1–11. 10.1007/s00401-011-0910-3

13. Cairns NJ, Bigio EH, Mackenzie IRA, et al (2007) Neuropathologic diagnostic and nosologic criteria for frontotemporal lobar degeneration: consensus of the Consortium for Frontotemporal Lobar Degeneration. Acta Neuropathol 114:5–22. 10.1007/s00401-007-0237-2

14. Hulsen T, de Vlieg J, Alkema W (2008) BioVenn – a web application for the comparison and visualization of biological lists using area-proportional Venn diagrams. BMC Genomics 9:488. 10.1186/1471-2164-9-488

15. Meneses A, Koga S, O’Leary J, et al (2021) TDP-43 Pathology in Alzheimer’s Disease. Mol Neurodegener 16:84. 10.1186/s13024-021-00503-x

16. Mann DMA, Snowden JS (2017) Frontotemporal lobar degeneration: Pathogenesis, pathology and pathways to phenotype. Brain Pathol 27:723–736. 10.1111/bpa.12486

17. Josephs KA, Murray ME, Tosakulwong N, et al (2019) Pathological, imaging and genetic characteristics support the existence of distinct TDP-43 types in non-FTLD brains. Acta Neuropathol 137:227–238. 10.1007/s00401-018-1951-7

18. Kawakami I, Arai T, Hasegawa M (2019) The basis of clinicopathological heterogeneity in TDP-43 proteinopathy. Acta Neuropathol 138:751–770. 10.1007/s00401-019-02077-x

19. Giannini LAA, Peterson C, Ohm D, et al (2021) Frontotemporal lobar degeneration proteinopathies have disparate microscopic patterns of white and grey matter pathology. Acta Neuropathol Commun 9:30. 10.1186/s40478-021-01129-2

20. Mol MO, Miedema SSM, van Swieten JC, et al (2021) Molecular Pathways Involved in Frontotemporal Lobar Degeneration with TDP-43 Proteinopathy: What Can We Learn from Proteomics? Int J Mol Sci 22:. 10.3390/ijms221910298

21. Mol MO, van Rooij JGJ, Wong TH, et al (2021) Underlying genetic variation in familial frontotemporal dementia: sequencing of 198 patients. Neurobiol Aging 97:148.e9-148.e16. 10.1016/j.neurobiolaging.2020.07.014

22. Hodges JR, Piguet O (2018) Progress and Challenges in Frontotemporal Dementia Research: A 20-Year Review. J Alzheimers Dis 62:1467–1480. 10.3233/JAD-171087

23. Shiarli AM, Jennings R, Shi J, et al (2006) Comparison of extent of tau pathology in patients with frontotemporal dementia with Parkinsonism linked to chromosome 17 (FTDP-17), frontotemporal lobar degeneration with Pick bodies and early onset Alzheimer’s disease. Neuropathol Appl Neurobiol 32:374–387. 10.1111/j.1365-2990.2006.00736.x

24. Bridel C, van Gils JHM, Miedema SSM, et al (2023) Clusters of co-abundant proteins in the brain cortex associated with fronto-temporal lobar degeneration. Alzheimers Res Ther 15:59. 10.1186/s13195-023-01200-1

25. Lemke N, Melis V, Lauer D, et al (2020) Differential compartmental processing and phosphorylation of pathogenic human tau and native mouse tau in the line 66 model of frontotemporal dementia. J Biol Chem 295:18508–18523. 10.1074/jbc.RA120.014890

26. Sanchez-Mejias E, Navarro V, Jimenez S, et al (2016) Soluble phospho-tau from Alzheimer’s disease hippocampus drives microglial degeneration. Acta Neuropathol 132:897–916. 10.1007/s00401-016-1630-5

27. Bulk M, Abdelmoula WM, Nabuurs RJA, et al (2018) Postmortem MRI and histology demonstrate differential iron accumulation and cortical myelin organization in early- and late-onset Alzheimer’s disease. Neurobiol Aging 62:231–242. 10.1016/j.neurobiolaging.2017.10.017

28. Swanson MEV, Scotter EL, Smyth LCD, et al (2020) Identification of a dysfunctional microglial population in human Alzheimer’s disease cortex using novel single-cell histology image analysis. Acta Neuropathol Commun 8:170. 10.1186/s40478-020-01047-9

29. Lee IH (2019) Mechanisms and disease implications of sirtuin-mediated autophagic regulation. Exp Mol Med 51:1–11. 10.1038/s12276-019-0302-7

30. Hinman JD, Chen C-D, Oh S-Y, et al (2008) Age-dependent accumulation of ubiquitinated 2’,3’-cyclic nucleotide 3’-phosphodiesterase in myelin lipid rafts. Glia 56:118–133. 10.1002/glia.20595

31. Warmus BA, Sekar DR, McCutchen E, et al (2014) Tau-mediated NMDA receptor impairment underlies dysfunction of a selectively vulnerable network in a mouse model of frontotemporal dementia. J Neurosci 34:16482–16495. 10.1523/JNEUROSCI.3418-14.2014

32. Ortiz-Sanz C, Balantzategi U, Quintela-López T, et al (2022) Amyloid β / PKC-dependent alterations in NMDA receptor composition are detected in early stages of Alzheimerś disease. Cell Death Dis 13:253. 10.1038/s41419-022-04687-y

33. Tracy TE, Madero-Pérez J, Swaney DL, et al (2022) Tau interactome maps synaptic and mitochondrial processes associated with neurodegeneration. Cell 185:712-728.e14. 10.1016/j.cell.2021.12.041

34. Toomey CE, Heywood WE, Evans JR, et al (2022) Mitochondrial dysfunction is a key pathological driver of early stage Parkinson’s. Acta Neuropathol Commun 10:134. 10.1186/s40478-022-01424-6

35. Tracy TE, Madero-Pérez J, Swaney DL, et al (2022) Tau interactome maps synaptic and mitochondrial processes associated with neurodegeneration. Cell 185:712-728.e14. 10.1016/j.cell.2021.12.041

36. Thal DR, Rüb U, Orantes M, Braak H (2002) Phases of A beta-deposition in the human brain and its relevance for the development of AD. Neurology 58:1791–1800. 10.1212/wnl.58.12.1791

37. Mirra SS, Heyman A, McKeel D, et al (1991) The Consortium to Establish a Registry for Alzheimer’s Disease (CERAD). Part II. Standardization of the neuropathologic assessment of Alzheimer’s disease. Neurology 41:479–486. 10.1212/wnl.41.4.479

38. Prokop S, Miller KR, Labra SR, et al (2019) Impact of TREM2 risk variants on brain region-specific immune activation and plaque microenvironment in Alzheimer’s disease patient brain samples. Acta Neuropathol 138:613–630. 10.1007/s00401-019-02048-2

39. Josephs KA, Hodges JR, Snowden JS, et al (2011) Neuropathological background of phenotypical variability in frontotemporal dementia. Acta Neuropathol 122:137–153. 10.1007/s00401-011-0839-6

40. Gorno-Tempini ML, Dronkers NF, Rankin KP, et al (2004) Cognition and anatomy in three variants of primary progressive aphasia. Ann Neurol 55:335–346. 10.1002/ana.10825

41. Landin-Romero R, Kumfor F, Leyton CE, et al (2017) Disease-specific patterns of cortical and subcortical degeneration in a longitudinal study of Alzheimer’s disease and behavioural-variant frontotemporal dementia. Neuroimage 151:72–80. 10.1016/j.neuroimage.2016.03.032

42. Hornberger M, Shelley BP, Kipps CM, et al (2009) Can progressive and non-progressive behavioural variant frontotemporal dementia be distinguished at presentation? J Neurol Neurosurg Psychiatr 80:591–593. 10.1136/jnnp.2008.163873

43. Walker JM, Kazempour Dehkordi S, Fracassi A, et al (2022) Differential protein expression in the hippocampi of resilient individuals identified by digital spatial profiling. Acta Neuropathol Commun 10:23. 10.1186/s40478-022-01324-9

