## Supplemental Figures for "High-plex Digital Spatial Profiling Identifies Subregion-Dependent Directed Proteome Changes Across Multiple Variants of Dementia"

**Supplemental Table 1**

Individual Protein Count significance and Fold Change in the White Matter of the Cortex and compared to NHC. NS represents no significant change when compared to NHC. A ↑ indicates a positive fold change and a ↓ indicates a negative fold change when comparing the disease group to NHC.

| | AD | c9ALS | FTLD-17 | FTLD-TDP | GRN |
|---|---|---|---|---|---|
| AB 42 | NS | ↓ p = 0.011, FC = 0.557 | NS | NS | NS |
| ApoA1 | NS | ↓ p = 0.025, FC = 0.363 | ↓ p = 0.001, FC = 0.209 | NS | NS |
| BDNF | NS | NS | ↑ p = 0.011, FC = 2.147 | NS | NS |
| Calbindin | NS | NS | ↓ p = 0.028, FC = 0.351 | NS | NS |
| CD11b | NS | NS | ↓ p = 0.022, FC = 0.463 | NS | NS |
| CNPase | NS | NS | ↓ p = 0.002, FC = 0.124 | NS | NS |
| FoxA2 | ↓ p = 0.032, FC = 0.488 | NS | NS | NS | NS |
| FUS | NS | NS | ↓ p = 0.036, FC = 0.511 | NS | NS |
| GAPDH | NS | ↓ p = 0.034, FC= 0.366 | NS | NS | NS |
| IBA1 | NS | NS | ↓ p = 0.017, FC = 0.285 | NS | NS |
| Lef1 | ↓ p =0.014, FC= 0.562 | NS | ↓ p = 0.019, FC = 0.579 | NS | NS |
| MAP2 | NS | NS | ↓ p = 0.034, FC = 0.189 | ↓ p =0.020, FC = 0.159 | NS |
| Ms.IgG2a | ↓ p =0.009, FC= 0.534 | NS | NS | NS | NS |
| NEFL | NS | ↓ p = 0.021, FC = 0.409 | NS | NS | NS |
| NMDAR1 | NS | NS | ↓ p = 0.029, FC= 0.353 | NS | NS |
| NRGN | NS | ↓ p = 0.006, FC = 0.203 | NS | NS | NS |
| Olig2 | NS | NS | ↓ p = 0.027, FC = 0.432 | NS | NS |
| P2ry12 | NS | NS | ↓ p = 0.005, FC = 0.210 | NS | NS |
| p53 | ↓ p = 0.048, FC =0.626 | NS | ↓ p = 0.016, FC = 0.567 | NS | NS |
| Park5 | NS | ↓ p = 0.026, FC= 0.362 | ↓ p <0.001, FC.= 0.154 | NS | ↓ p = 0.009, FC – 0.298 |
| polyGP | ↓ p = 0.017, FC = 0.561 | ↓ p = 0.009, FC = 0.553 | ↓ p = 0.043, FC = 0.636 | NS | NS |
| PSEN1 | NS | NS | NS | ↓ p = 0.034, FC = 0.457 | NS |
| Phospho. Tau S214 | NS | NS | ↓ p = 0.003, FC = 5.946 | NS | ↓ p = 0.043, FC = 3.206 |
| Phospho. Tau S404 | ↓ p = 0.046, FC = 0.422 | NS | ↓ p = 0.009, FC = 0.313 | NS | NS |
| Phospho. Tau T231 | NS | ↓ p = 0.042, FC = 0.441 | NS | NS | NS |
| SIRT2 | NS | NS | ↓ p = 0.006, FC = 0.344 | NS | NS |
| SNCA_filament | NS | NS | ↓ p = 0.036, FC = 0.385 | NS | NS |
| Tau | NS | NS | ↓ p = 0.009, FC= 0.263 | NS | NS |
| TDP43 | NS | NS | ↓ p = 0.032, FC = 0.546 | NS | NS |
| TMEM119 | NS | NS | ↓ p = 0.004, FC = 0.151 | NS | NS |
| Tuj1 | NS | NS | ↓ p = 0.002, FC = 0.334 | NS | NS |
| UBB | NS | NS | ↓ p = 0.008. FC = 0.257 | NS | ↓ p = 0.043, FC = 0.362 |

**Supplemental Table 2**

Individual Protein Count significance and Fold Change in cortical layer II-V of the compared to NHC. NS represents no significant change when compared to NHC. A ↑ indicates a positive fold change and a ↓ indicates a negative fold change when comparing the disease group to NHC.

| | AD | c9ALS | FTLD-17 | FTLD-TDP | GRN |
|---|---|---|---|---|---|
| AB 42 | ↑ p = 0.003, FC = 2.566 | NS | NS | NS | NS |
| APP | NS | NS | ↓ p = 0.024, FC = 0.294 | ↓ p = 0.016, FC = 0.0271 | NS |
| BACE1 | NS | NS | ↓ p = 0.033, FC = 0.325 | NS | NS |
| Calbindin | NS | NS | ↓ p = 0.021, FC = 0.295 | NS | NS |
| CNPase | NS | NS | ↓ p = 0.019, FC = 0.279 | ↓ p = 0.050, FC = 0.348 | NS |
| FUS | NS | NS | ↓ p = 0.020, FC – 0.401 | NS | NS |
| GAPDH | NS | NS | ↓ p = 0.004, FC = 0.386 | NS | NS |
| GPNMB | NS | NS | ↓ p = 0.036, FC = 0.469 | NS | NS |
| HISTONE H3 | NS | NS | ↓ p = 0.037. FC = 0.216 | NS | NS |
| IBA1 | NS | NS | ↓. p = 0.007, FC 0.222 | NS | NS |
| IQGAP2 | NS | NS | ↓ p = 0.023, FC = 0.469 | NS | NS |
| Ki67 | NS | NS | NS | ↓ p=0.045, FC = 0.440 | NS |
| MAP2 | NS | NS | NS | NS | NS |
| NEFL | NS | ↓ p = 0.041, FC = 0.413 | NS | NS | ↓ p = 0.039, FC = 0.409 |
| NMDAR1 | NS | NS | NS | ↓ p = 0..047, FC = 0.301 | NS |
| NRGN | NS | ↓ p = 0.026, FC = 0.296 | NS | NS | NS |
| Olig2 | NS | NS | ↓ p = 0.018, FC = 0.382 | ↓ p = 0.032, FC = 0.420 | |
| P2ry12 | ↓ p = 0.015, FC = 0.402 | ↓ p = 0.010, FC = 0.379 | ↓ p =0.002, FC = 0.291 | ↓ p = 0.012, FC =0.389 | ↓ p = 0.012, FC = 0.390 |
| p53 | NS | NS | ↓ p = 0.040, FC = 0.475 | NS | |
| Park5 | NS | NS | ↓ p <0.001, FC = 0.191 | NS | ↓ p = 0.019, FC = 0.340 |
| Phospho. Tdp. 43 | NS | NS | NS | NS | ↓ p = 0.016, FC = 4.004 |
| PINK1 | NS | ↓ p= 0.044, FC 0.440 | NS | NS | NS |
| Phospho. Tau S214 | ↑ p <0.001, FC = 12.596 | NS | ↑ p <0.001, FC = 4.949 | NS | NS |
| Phospho. Tau S396 | ↑ p <0.001, FC = 11.277 | NS | NS | NS | NS |
| Phospho. Tau 231 | ↑ p = 0.004, FC = 4.467 | NS | NS | NS | NS |
| S6 | NS | NS | ↓ p = 0.008, FC = 0.238 | ↓ p = 0.046 FC = 0.351 | NS |
| SIRT2 | NS | NS | ↓ p = 0.011, FC = 0.346 | NS | NS |
| Tau | NS | NS | ↓ p =0.044, FC = 0.348 | NS | NS |
| TDP 43 | NS | NS | ↓ p = 0.010. FC = 0.408 | ↓ p=0.016, FC =0.436 | NS |
| TH | NS | NS | ↓ p.= 0.024, FC = 0.459 | NS | NS |
| TMEM119 | NS | NS | ↓ p = 0.006, FC = 0.208 | NS | NS |
| Tuj1 | NS | NS | ↓ p =0.020, FC = 0.451 | NS | NS |
| UBB | NS | NS | NS | ↓ p = 0.036, FC = 0.340 | NS |
| vGlut1 | NS | NS | NS | ↓ p = 0.042, FC = 0.322 | NS |

**Supplemental Table 3**

Individual Protein Count significance and Fold Change in cortical layer I compared to NHC. NS represents no significant change when compared to NHC. A ↑ indicates a positive fold change and a ↓ indicates a negative fold change when comparing the disease group to NHC.

| | AD | c9ALS | FTLD-17 | FTLD-TDP | GRN |
|---|---|---|---|---|---|
| AB 42 | ↑ p = 0.002, FC = 3.261 | NS | NS | NS | NS |
| APP | NS | NS | NS | ↓ p = 0.038, FC = 0.285 | NS |
| CNPase | NS | NS | ↓ p = 0.026, FC = 0.238 | NS | NS |
| GAPDH | NS | ↓ p = 0.036, FC = 0.400 | NS | NS | NS |
| IBA1 | NS | NS | ↓ p = 0.021, FC = 0.316 | NS | NS |
| NMDAR1 | NS | ↓ p = 0.007, FC = 0.160 | ↓ **p = 0.039, FC = 0.236** | ↓ p = 0.047, FC = 0.236 | ↓ p = 0.025, FC = 0.221 |
| NRGN | NS | ↓ p= 0.018, FC = 0.281 | NS | NS | NS |
| Park5 | NS | NS | ↓ p = 0.025, FC = 0.418 | NS | NS |
| Phospho. Tdp. 43 | NS | ↑ p= 0.026, FC = 4.511 | NS | NS | ↑ p = 0.029, FC = 4.387 |
| Phospho. Tau S214 | ↑ p <0.001, FC = 11.825 | NS | ↑ p = 0.003, FC – 4.650 | NS | NS |
| Phospho. Tau S396 | ↑ p < 0.001, FC =13.871 | NS | ↑ p = 0.075, FC = 2.905 | NS | NS |
| Phospho. Tau T231 | ↑ p < 0.001, FC = 6.426 | NS | NS | NS | NS |
| S100B | NS | ↓ p = 0.045, FC = 0.494 | NS | NS | NS |
| SIRT2 | NS | NS | ↓ p = 0.002, FC 0.111 | NS | ↓ p = 0.039, FC = 0.270 |
| Tuj1 | NS | NS | NS | NS | NS |

**Supplemental Table 4**

Individual Protein Count significance and Fold Change when all ROIs are combined and compared to NHC. NS represents no significant change when compared to NHC. A ↑ indicates a positive fold change and a ↓ indicates a negative fold change when comparing the disease group to NHC.

| | AD | c9ALS | FTLD-17 | FTLD-TDP | GRN |
|---|---|---|---|---|---|
| AB 42 | ↑ p = 0.003, FC = 2.566 | NS | NS | NS | NS |
| APP | NS | NS | ↓ p = 0.024, FC = 0.294 | ↓ p = 0.016, FC = 0.0271 | NS |
| BACE1 | NS | NS | ↓ p = 0.033, FC = 0.325 | NS | NS |
| Calbindin | NS | NS | ↓ p = 0.021, FC = 0.295 | NS | NS |
| CNPase | NS | NS | ↓ p = 0.019, FC = 0.279 | ↓ p = 0.050, FC = 0.348 | NS |
| FUS | NS | NS | ↓ p = 0.020, FC – 0.401 | NS | NS |
| GAPDH | NS | NS | ↓ p = 0.004, FC = 0.386 | NS | NS |
| GPNMB | NS | NS | ↓ p = 0.036, FC = 0.469 | NS | NS |
| HISTONE H3 | NS | NS | ↓ p = 0.037. FC = 0.216 | NS | NS |
| IBA1 | NS | NS | ↓. p = 0.007, FC 0.222 | NS | NS |
| IQGAP2 | NS | NS | ↓ p = 0.023, FC = 0.469 | NS | NS |
| Ki67 | NS | NS | NS | ↓ p=0.045, FC = 0.440 | NS |
| MAP2 | NS | NS | NS | NS | NS |
| NEFL | NS | ↓ p = 0.041, FC = 0.413 | NS | NS | ↓ p = 0.039, FC = 0.409 |
| NMDAR1 | NS | NS | NS | ↓ p = 0..047, FC = 0.301 | NS |
| NRGN | NS | ↓ p = 0.026, FC = 0.296 | NS | NS | NS |
| Olig2 | NS | NS | ↓ p = 0.018, FC = 0.382 | ↓ p = 0.032, FC = 0.420 | |
| P2ry12 | ↓ p = 0.015, FC = 0.402 | ↓ p = 0.010, FC = 0.379 | ↓ p =0.002, FC = 0.291 | ↓ p = 0.012, FC =0.389 | ↓ p = 0.012, FC = 0.390 |
| p53 | NS | NS | ↓ p = 0.040, FC = 0.475 | NS | |
| Park5 | NS | NS | ↓ p <0.001, FC = 0.191 | NS | ↓ p = 0.019, FC = 0.340 |
| Phospho. Tdp. 43 | NS | NS | NS | NS | ↓ p = 0.016, FC = 4.004 |
| PINK1 | NS | ↓ p= 0.044, FC 0.440 | NS | NS | NS |
| Phospho. Tau S214 | ↑ p <0.001, FC = 12.596 | NS | ↑ p <0.001, FC = 4.949 | NS | NS |
| Phospho. Tau S396 | ↑ p <0.001, FC = 11.277 | NS | NS | NS | NS |
| Phospho. Tau 231 | ↑ p = 0.004, FC = 4.467 | NS | NS | NS | NS |
| S6 | NS | NS | ↓ p = 0.008, FC = 0.238 | ↓ p = 0.046 FC = 0.351 | NS |
| SIRT2 | NS | NS | ↓ p = 0.011, FC = 0.346 | NS | NS |
| Tau | NS | NS | ↓ p =0.044, FC = 0.348 | NS | NS |
| TDP 43 | NS | NS | ↓ p = 0.010. FC = 0.408 | ↓ p=0.016, FC =0.436 | NS |
| TH | NS | NS | ↓ p.= 0.024, FC = 0.459 | NS | NS |
| TMEM119 | NS | NS | ↓ p = 0.006, FC = 0.208 | NS | NS |
| Tuj1 | NS | NS | ↓ p =0.020, FC = 0.451 | NS | NS |
| UBB | NS | NS | NS | ↓ p = 0.036, FC = 0.340 | NS |
| vGlut1 | NS | NS | NS | ↓ p = 0.042, FC = 0.322 | NS |

### **Supplementary Table 5 .** Individual case demographics and cognitive status

| Case | Disease | Primary Neuropathologic Diagnosis | Other Neuropathologic Diagnosis - AD | Other Neuropathologic Diagnosis - LBD | Other Neuropathologic Diagnosis - TDP | Other Neuropathologic Diagnosis - tau | Braak Stage | ABC | CERAD | Thal | Age at Onset | Age at Death | Duration (years) | ApoE | Race | Sex | Clin Dx | Family Hx |
|---|---|---|---|---|---|---|---|---|---|---|---|---|---|---|---|---|---|---|
| 1 | FTLD-TDP | FTLD-TDP | | | | | I | 0 | 0 | 0 | 55 | 61 | 6 | E3/3 | w | m | FTD, MND/ALS | Yes |
| 2 | FTLD-TDP | FTLD-TDP | | | | | I | 1 | 0 | 1 | 56 | 64 | 8 | E3/4 | w | f | FTD | Yes |
| 3 | AD | AD | | LBD-amygdala | | | VI | 3 | 3 | 5 | 56 | 64 | 8 | E3/4 | w | m | AD | |
| 4 | AD | AD | | LBD-amygdala | | | VI | 3 | 3 | 5 | 59 | 72 | 14 | E3/4 | w | f | AD | Yes |
| 5 | Control | AD | | LBD-amygdala | | | VI | 3 | 3 | 5 | 56 | 64 | 8 | E3/4 | w | m | AD | |
| 6 | AD | AD | | LBD-amygdala | LATE-NC (stage 1) | | VI | 3 | 3 | 5 | 51 | 64 | 13 | E3/4 | w | f | AD | na |
| 7 | GRN | FTLD-TDP (GRN mutation) | | | | | 0 | 0 | 0 | 0 | 52 | 62 | 10 | E3/3 | other | f | FTD | Yes |
| 8 | FTLD-TDP | FTLD-TDP | | | | | II | 0 | 0 | 0 | 62 | 71 | 9 | E3/3 | w | f | FTD | No |
| 9 | AD | AD | | | LATE-NC (stage 1) | | V | 3 | 3 | 5 | 52 | 62 | 10 | E3/4 | w | m | AD | No |
| 10 | Control | Control | | | | | I | 0 | 0 | 0 | na | 59 | | E2/3 | b | m | ischemic heart disease and fibrosis | |
| 11 | AD | AD | | | LATE-NC (stage 2) | | VI | 3 | 3 | 5 | 52 | 60 | 8 | E4/4 | b | m | Pick's disease | Yes |
| 12 | c9ALS | FTLD-TDP (C9 expansion) | | | | | III | 1 | 0 | 2 | 57 | 66 | 9 | E3/3 | w | m | FTD/Pick's disease | Yes |
| 13 | GRN | FTLD-TDP (GRN mutation) | | | LATE-NC (stage 3) | | I | 1 | 0 | 2 | 67 | 71 | 4.5 | E3/4 | w | m | AD | Yes |
| 14 | AD | AD | | | | | VI | 3 | 3 | 5 | 53 | 58 | 5 | E3/4 | w | f | AD | No |
| 15 | FTLD-TDP | FTLD-TDP | AD | | | | IV | 2 | 3 | 5 | 58 | 60 | 2 | E3/4 | w | f | FTD | Yes |
| 16 | FTLD-17 | FTDP-17 (P301L) | | | | | na | 0 | 0 | 0 | 56 | 60 | 4 | E3/3 | w | f | FTDP-17 (P301L) | Yes |
| 17 | FTLD-17 | FTDP-17 (P301L) | | | | | 0 | 0 | 0 | 0 | 56 | 64 | 8.5 | E3/4 | w | m | FTDP-17 (P301L) | Yes |
| 18 | FTLD-TDP | FTLD-TDP | | | | | II | 1 | 0 | 1 | 66 | 67 | 1 | E2/3 | w | m | FTD; CBD vs. prion disease | Yes |
| 19 | GRN | FTLD-TDP (GRN mutation) | | LBD-amygdala | | | I | 1 | 0 | 2 | 58 | 62 | 4.5 | E3/4 | w | m | CBD | Yes |
| 20 | GRN | FTLD-TDP (GRN mutation) | | | | | I | 0 | 0 | 0 | 57 | 63 | 6 | E2/3 | w | m | primary progressive aphasia | No |
| 21 | c9ALS | FTLD-TDP (C9 expansion) | | | ALS (C9 expansion) | | 0 | 0 | 0 | 0 | 55 | 57 | 1.5 | na | w | f | ALS+FTD | na |
| 22 | c9ALS | FTLD-TDP (C9 expansion) | | LBD-amygdala | | | IV | 1 | 0 | 1 | 60 | 70 | 10 | E2/3 | w | f | FTD-primary progressive aphasia; PSP | No |
| 23 | GRN | FTLD-TDP (GRN mutation) | | | | | III | 1 | 0 | 1 | 55 | 61 | 6 | E3/3 | w | f | FTD | Yes |
| 24 | Control | Control | | | | | II | 1 | 0 | 1 | na | 61 | | na | b | f | diabetes; hx hip replacement | |
| 25 | c9ALS | FTLD-TDP (C9 expansion) | | | ALS | | II | 0 | 0 | 0 | 62 | 66 | 4 | E4/4 | w | m | FTD; ALS | No |
| 26 | c9ALS | FTLD-TDP (C9 expansion) | | | ALS | Tau pathology-mild | na | na | 0 | 1 | 58 | 67 | 9 | | w | m | FTD; ALS | Yes |
| 27 | FTLD-17 | FTDP-17 (P301L) | | | | | na | 0 | 0 | 0 | 51 | 56 | 5 | na | w | f | FTD (probable FTDP-17) | Yes |
| 28 | c9ALS | FTLD-TDP (C9 expansion) | | | TDP-43 incl in sp cord neurons | | III | 1 | 0 | 1 | 56 | 7 | 14 | E3/3 | w | f | FTD | Yes |
| 29 | GRN | FTLD-TDP (GRN mutation) | | | | | I | 1 | 0 | 2 | 55 | 61 | 6 | E3/4 | w | f | FTD (GRN mutation) | Yes |
| 30 | Control | Control | | | | | I | 1 | 0 | 2 | na | 70 | | E3/3 | b | m | control | Yes |
| 31 | FTLD-17 | FTDP-17 (P301L presumed) | AD-probable | | | | na | na | 2 | 4 | 56 | 65 | 9 | E2/3 | w | f | FTD | Yes |
| 32 | FTLD-17 | FTDP-17 (G389R) | | | | | na | na | 0 | 1 | 36 | 40 | 4 | E3/3 | w | m | FTD | Yes |
| 33 | Control | Control | | | | | I | 0 | 0 | 0 | na | 65 | na | E3/3 | w | f | systemic primary amyloidosis, AL type | na |
| 34 | Control | Control | | | | | II | 1 | 0 | 2 | na | 61 | | E3/4 | b | m | diabetes, renal disease, congestive heart failure | |
| 35 | FTLD-TDP | FTLD-TDP | | LBD-amygdala | | | 0 | 0 | 0 | 0 | 52 | 59 | 7 | E3/3 | w | m | FTD | Yes |
| 36 | FTLD-17 | FTDP-17 (R406W) | AD-definite | | | | na | na | 2 | 3 | na | 65 | na | na | w | f | FTDP-17 (R406W) | Yes |
